# Real-Time Tracking of Selective Auditory Attention from M/EEG: A Bayesian Filtering Approach

**DOI:** 10.1101/222661

**Authors:** Sina Miran, Sahar Akram, Alireza Sheikhattar, Jonathan Z. Simon, Tao Zhang, Behtash Babadi

## Abstract

Humans are able to identify and track a target speaker amid a cacophony of acoustic interference, which is often referred to as the cocktail party phenomenon. Results from several decades of studying this phenomenon have culminated in recent years in various promising attempts to decode the attentional state of a listener in a competing-speaker environment from non-invasive neuroimaging recordings such as magnetoencephalography (MEG) and electroencephalography (EEG). To this end, most existing approaches compute correlation-based measures by either regressing the features of each speech stream to the M/EEG channels (the decoding approach) or vice versa (the encoding approach). These procedures operate in an offline fashion, i.e., require the entire duration of the experiment and multiple trials to provide robust results. Therefore, they cannot be used in emerging applications such as smart hearing aid devices, where a single trial must be used in real-time to decode the attentional state. In this paper, we close this gap by developing an algorithmic pipeline for real-time decoding of the attentional state. Our proposed framework consists of three main modules: 1) Real-time and robust estimation of encoding or decoding coefficients, achieved by sparse adaptive filtering, 2) Extracting reliable markers of the attentional state, and thereby generalizing the widely-used correlation-based measures thereof, and 3) Devising a near real-time state-space estimator that translates the noisy and variable attention markers to robust and reliable estimates of the attentional state with minimal delay. Our proposed algorithms integrate various techniques including forgetting factor-based adaptive filtering, ℓ_1_-regularization, forward-backward splitting algorithms, fixed-lag smoothing, and Expectation Maximization. We validate the performance of our proposed framework using comprehensive simulations as well as application to experimentally acquired M/EEG data. Our results reveal that the proposed real-time algorithms perform nearly as accurate as the existing state-of-the-art offline techniques, while providing a significant degree of adaptivity, statistical robustness, and computational savings.

## 1. Introduction

The ability to select a single speaker in an auditory scene, consisting of multiple competing speakers, and maintain attention to that speaker is one of the hallmarks of human brain function. This phenomenon has been referred to as the cocktail party effect [1, 2, 3]. The mechanisms underlying the real-time process by which the brain segregates multiple sources in a cocktail party setting has been the topic of active research for decades [4, 5]. Although the details of these mechanisms are for the most part unknown, various studies have underpinned the role of specific neural processes involved in this function. As the acoustic signals propagate through the auditory pathway, they are decomposed into spectrotemporal features at different stages, and a rich representation of the complex auditory environment reaches the auditory cortex. It has been hypothesized that the perception of an auditory object is the result of adaptive binding as well as discounting of these features [6, 7, 8, 9].

From a computational modeling perspective, there have been several attempts at designing so-called “attention decoders”, where the goal is to reliably decode the attentional focus of a listener in a multi-speaker environment using non-invasive neuroimaging techniques like electroencephalography (EEG) [10, 11, 12] and magnetoencephalography (MEG) [13, 14, 15, 16, 17]. These methods are typically based on reverse correlation or estimating linear encoding/decoding models using off-line regression techniques, and thereby detecting salient peaks in the model coefficients that are modulated by the attentional state [18]. The aforementioned salient peaks have been observed at a typical lag of ~ 200 ms for EEG [11] and ~ 100 ms for MEG [13], implying the longer-lasting effect and further processing of the attended stimuli as compared to the unattended ones.

Although the foregoing approaches have proven successful in reliable attention decoding, they have two major limitations that make them unsuitable for emerging real-time applications such as Brain-Computer Interface (BCI) systems and smart hearing aids. First, the temporal resolution for decoding the attentional state is on the order of tens of seconds, whereas humans can switch their attention from one speaker to another at a much shorter time scale. This is due to their so-called “batch-mode” design, which requires the entire data from one or multiple trials at once for processing. Second, approaches based on linear regression (e.g., reverse correlation) need large training datasets, often from multiple subjects and trials, to estimate the decoder/encoder reliably. Access to such training data is only possible through repeated calibration stages, which may not always be possible in real-time applications. While recent results [15, 16] address the first shortcoming by employing state-space models and thereby producing robust estimates of the attentional state from limited data, they are not yet suitable for real-time applications.

In this paper, we close this gap by designing a modular framework for real-time attention decoding from non-invasive M/EEG recordings that overcomes the aforementioned limitations using techniques from Bayesian filtering. Our proposed framework includes three main modules. The first module pertains to estimating *dynamic* models of decoding/encoding in *real-time*. To this end, we use the forgetting factor mechanism of the Recursive Least Squares (RLS) algorithm together with the ℓ_1_ regularization penalty from Lasso to capture the dynamics in the data while preventing overfitting [17, 19]. The real-time inference is then efficiently carried out using a Forward-Backward Splitting (FBS) procedure [20]. In the second module, we extract an attention-modulated feature, which we refer to as “attention marker”, as a function of the M/EEG recordings, the estimated encoding/decoding coefficients, and the auditory stimuli. For instance, the attention marker can be a correlation-based measure or the magnitude of certain peaks in the model coefficients. We carefully design the attention marker features to capture the attention modulation and thereby maximally separate the contributions of the attended and unattended speakers in the neural response in both MEG and EEG applications.

The extracted features are then passed to a novel state-space estimator in the third module, and thereby are translated into robust and dynamic measures of the attentional state. The state-space estimator is based on Bayesian fixed-lag smoothing, and operates in *near real-time* with controllable delay. The fixed-lag design creates a trade-off between real-time operation and robustness to stochastic fluctuations. In addition, we modify the Expectation-Maximization algorithm and the nonlinear filtering and smoothing techniques of [16] for real-time implementation. Compared to existing techniques, our algorithms require minimal supervised data for initialization and tuning. In order to validate our real-time attention decoding algorithms, we apply them to both simulated and experimentally recorded EEG and MEG data in dual-speaker environments. Our results suggest that the performance of our proposed framework is comparable to the state-of-the-art batch-mode algorithms of [10, 12, 16], while operating in near real-time with ~ 1s delay.

The rest of the paper is organized as follows: In Section 2, we develop the three main modules in our proposed framework as well as the corresponding estimation algorithms. We present the application of our framework to both synthetic and experimentally recorded M/EEG data in Section 3, followed by discussion and concluding remarks in Section 4.

## 2. Material and Methods

Figure 1 summarizes our proposed framework for real-time tracking of selective auditory attention from M/EEG. In the *Dynamic Encoder/Decoder Estimation* module, the encoding/decoding models are fit to neural data in real-time. The *Attention Marker* module uses the estimated model coefficients as well as the recorded data to compute a feature that is modulated by the instantaneous attentional state. Finally, in the *State-Space Model* module, the foregoing features are refined through a linear state-space model with nonlinear observations, resulting in robust and dynamic estimates of the attentional state.

**Figure 1:**
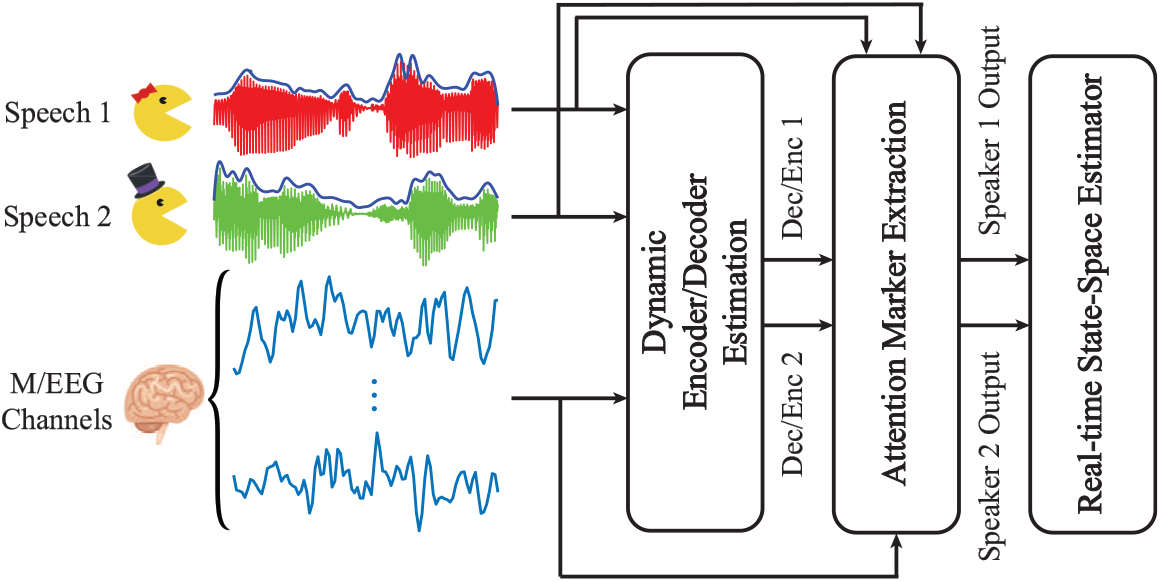
A schematic depiction of our proposed framework for real-time tracking of selective auditory attention from M/EEG.

In Section 2.1, we formally define the dynamic encoding and decoding models, and develop low-complexity and real-time techniques for their estimation. This is followed by Section 2.2, in which we define suitable attention markers for M/EEG inspired by existing literature. In Section 2.3, we propose a state-space model that processes the extracted attention markers in order to produce near real-time estimates of the attentional state with minimal delay.

### 2.1. Dynamic Encoding and Decoding Models

The role of a neural encoding model is to map the stimulus to the neural response. Inspired by existing literature on attention decoding [13, 10, 16], we take the speech envelopes as covariates representing the stimuli. The neural response is manifested in the M/EEG recordings. Encoding models can be used to predict the neural response from the stimulus. In contrast, in a neural decoding model, the goal is to express the stimulus as a function of the neural response. Inspired by previous studies, we consider linear encoding and decoding models in this work.

The encoding and decoding models can be cast as mathematically dual formulations. In a dual-speaker environment, let 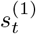 and 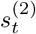 denote the speech envelopes (in logarithmic scale), corresponding to speakers 1 and 2, respectively, for *t* = 1, 2,…, *T*. Also, let 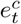 denote the neural response recorded at time *t* and channel *c*, for *c* =1, 2,…, *C*. Throughout the paper, we assume the same sampling frequency for both the M/EEG channels and the envelopes. Consider consecutive and non-overlapping windows of length *W*, and define 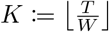. We consider piece-wise constant dynamics for the encoding and decoding coefficients, in which the coefficients assume to be constant over each window.

In the encoding setting, we define the vector 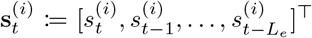 for *i* = 1, 2, where *L_e_* is the total lag considered in the model. Also, let *E_t_* denote a generic linear combination of 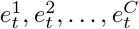 with some fixed set of weights. These weights can be set to select a single channel, i.e., 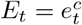 for some *c*, or they can be pre-estimated from training data so that *E_t_* represents the dominant auditory component of the neural response [21]. The encoding coefficients then relate 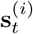 to *E_t_*. In the decoding setting, we define the vector 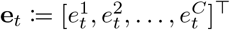 and 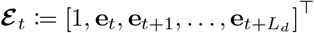, where *L_d_* is the total lag in the decoding model and determines the extent of future neural responses affected by the current stimuli. The decoding coefficients then relate 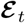 to 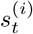.

Our goal is to recursively estimate the encoding/decoding coefficients in a real-time fashion as the new data samples become available. In addtion, we aim to simultaneously induce adaptivity of the parameter estimates and capture their sparsity. To this end, we employ the following generic optimization problem:

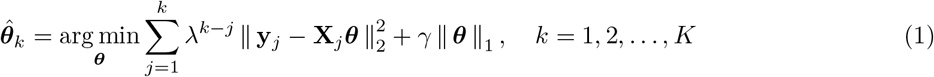

where **y**_*j*_ and **X**_*j*_ are the vector of response variables and the matrix of covariates pertinent to window *j*, ***θ*** is the parameter vector, λ ∈ (0,1] is the forgetting factor, and *γ* is a regularization parameter. The optimization problem of Eq. 1 is a modified version of the LASSO problem [22].

For the encoding problem, we define **y**_*k*_ := [*E*_(*k*−1)*W* + 1_; *E*_(*k*−1)*W*+2_;…; *E_kW_*]^⊤^ and 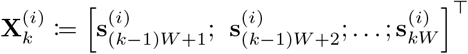, for *k* = 1, 2,…,*K* and *i* =1, 2. Therefore, the full encoding covariate matrix at the *k^th^* window is defined as 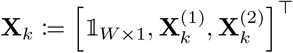, where the all-ones vector 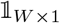 corresponds to the regression intercept. In the decoding problem, we define 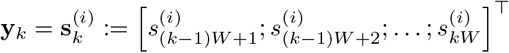, where *i* ∈ {1, 2}. Also, the full decoding covariate matrix at the *k^th^* window is 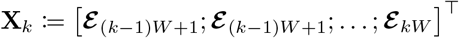, for *k* = 1, 2,…, *K*.

The optimization problem of Eq. (1) has a useful Bayesian interpretation: if the observation noise were i.i.d. Gaussian, and the parameters were exponentially distributed, it is akin to the maximum *a posteriori* (MAP) estimate of the parameters. The quadratic terms correspond to the exponentially-weighted log-likelihood of the observations up to window *k*, and the ℓ_1_-norm corresponds to the log-density of an independent exponential prior on the elements of ***θ***. The exponential prior serves as an effective regularization to promote sparsity of the estimate 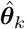. Note that we have 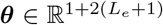 for the encoding model and 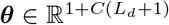 for the decoding model in (1).

*Remark* 1. The hyperparameter λ provides a tradeoff between the adaptivity and the robustness of estimated coefficients, and it can be determined based on the inherent dynamics in the data. The case of λ =1 corresponds to the natural data log-likelihood, i.e., the batch-mode parameter estimates. It has been shown that 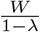 can serve as the *effective* number of recent samples used to calculate 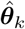 in (1) [23]. The parameter 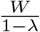 can also be viewed as the dynamic integration time: it needs to be chosen long enough so that the estimation is stable, but also short enough to be able to capture the dynamics of neural process involved in switching attention. The hyperparameter *γ* controls the tradeoff between the Maximum Likelihood (ML) fit and the sparsity of estimated coefficients, and it is usually determined through cross-validation.

*Remark* 2. In the decoding problem, Eq. (1) is solved separately at each window for each speech envelope, resulting in a set of decoding coefficients per speaker. In the encoding setting, we combine the stimuli as explained and solve Eq. (1) once at each window to obtain both of the encoder estimates. If the encoding/decoding coefficients are expected to be sparse in a basis represented by the columns of a matrix G, such as the Haar or Gabor bases, we can replace **X**_*j*_ in (1) by **X**_*j*_**G**, for *j* = 1, 2,…, *k*, and solve for 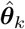 as before. Then, the final encoding/decoding coefficients are given by 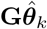. In the context of encoding models, the coefficients are referred to as the Temporal Response Function (TRF) [13, 17]. The TRFs are known to exhibit some degree of sparsity and smoothness in the lag domain, which can be represented over a basis consisting of shifted Gaussian kernels (see [17] for details).

*Remark* 3. Throughout the paper, we assume that the envelopes of the clean speeches are available. Given that this assumption does not hold in practical scenarios, recent algorithms on the extraction of speech envelopes from acoustic mixtures [24, 25, 26, 27, 28] can be added as a pre-processing module to our framework.

Among the many existing algorithms for solving the modified LASSO problem of Eq. (1), we choose the Forward-Backward Splitting (FBS) algorithm [20], also known as the proximal gradient method. When coupled with proper step-size adjustment methods, FBS is well-suited for real-time and low-complexity updates of 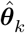 at each window. In this work, we have used the FASTA software package [29] available online [30], which has built-in features for all the FBS stepsize adjustment methods. A detailed overview of the FBS algorithm and its properties is given in Section S1 of the Supplementary Material.

### 2.2 Attention Markers

We define the *attention marker* as a mapping function from the estimated encoding/decoding coefficients for each speaker as well as the data in each window to positive real numbers. To be more precise, at window *k* and for speaker *i*, in the context of encoding models, the attention marker takes the speaker’s estimated encoding coefficients 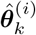 the speaker’s covariate matrix 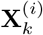, and the M/EEG responses **y**_*k*_ as inputs; similarly, in the context of decoding models, the attention marker takes the speaker’s estimated decoding coefficients 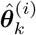, the M/EEG covariate matrix **X**_*k*_, and the speaker’s speech envelope vector 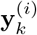 as inputs. In both cases, the attention marker outputs a positive real number, which we denote by 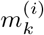 henceforth, for *i* = 1, 2 and *k* = 1, 2,…, *K*. Thus, in the modular design of Fig. 1, at each window *k*, the two outputs 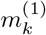 and 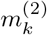 are passed from the Attention Maker module to the State-Space Model module as measures of the attentional state at window *k*.

In [10], a correlation-based measure has been adopted in the decoding model to classify the attended and the unattended speeches in a dual-speaker environment. The approach in [10] is based on estimating an *attended* decoder from the training data to reconstruct the speech envelope from EEG for each trial. Then, the correlation of this reconstructed envelope with each of the two speech envelopes is computed, and the speaker with the larger correlation coefficient is deemed as the attended speaker. This method cannot be directly applied to the real-time setting, since the lack of abundant training data hinders a reliable estimate of the *attended* decoder. However, assuming that the auditory M/EEG response is more influenced by the attended speaker than the unattended one, we can expect that the decoder corresponding to the *attended* speaker exhibits a higher performance in reconstructing the speech envelope it has been trained on, as suggested by the classification comparisons in [10]. Inspired by these results, we can define the attention marker in the decoding scenario as the correlation magnitude between the speech envelope and its reconstruction by the corresponding decoder, i.e., 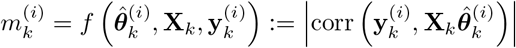 for *i* = 1, 2 and *k* = 1, 2,…, *K*. As we will demonstrate later in Section 3, this attention marker is suitable for the analysis of EEG recordings.

In the context of cocktail party studies using MEG, it has been shown that the magnitude of the negative peak in the TRF of the attended speaker around a lag of 100 ms, referred to as the M100 component, is higher than that of the unattended speaker [13, 17, 16]. Inspired by these findings, in the encoding scenario applied to MEG data, we can define the attention marker 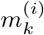 to be the magnitude of the 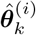 coefficients corresponding to the M100 component, for *i* = 1, 2 and *k* = 1, 2,…, *K*.

Due to the inherent uncertainties in the M/EEG recordings, the limitations of non-invasive neuroimaging in isolating the relevant neural processes, and the unknown and likely nonlinear processes involved in auditory attention, the foregoing attention markers derived from linear models are not readily reliable indicators of the attentional state. Given ample training data, however, these attention markers have been validated using batch-mode analysis. However, their usage in a real-time setting requires more care, as the limited data in real-time applications adds a major source of uncertainty to the foregoing list. To address this issue, a state-space model is required in the real-time setting to correct for the uncertainties and stochastic fluctuations of the attention markers caused by the limited integration time in real-time application. We will discuss in detail the formulation and advantages of such a state-space model in the following subsection.

### 2.3 State-Space Model

In order to translate the attention markers 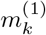 and 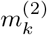, for *k* = 1, 2,…, *K*, into a robust and statistically interpretable measure of the attentional state, we employ state-space models. Inspired by the models used in [16], we design a new state-space model and a corresponding estimator that operates in a fixed-lag smoothing fashion, and thereby admits real-time processing while maintaining the benefits of batch-mode state-space models. Recall that the index *k* corresponds to a window in time ranging from *t* = (*k* − 1)*W* + 1 to *t* = *kW*; however, we refer to each index *k* as an *instance* when talking about the state-space model not to be confused with the sliding window of the fixed-lag design.

Figure 2 displays the fixed-lag smoothing design of the state-space estimator. Suppose that we are at the instance *k* = *k*_0_. We consider a window of length *K_W_* = *K_B_* + *K_F_* + 1 as shown in Fig. 2, where *K_F_* and *K_B_* are respectively called the forward-lag and the backward-lag. In order to carry out the computations in real-time, we assume all of the attentional state estimates to be fixed prior to this window and only update our estimates for the instances within, based on 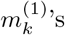 and 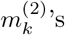 inside the window. In a fixed-lag framework, at *k* = *k*_0_, the goal is to provide an estimate of the attentional state at instance *k* = *k*\*, where *k*\* = *k*_0_–*K_F_*. The parameter *K_F_* creates a tradeoff between real-time and robust estimation of the attentional state. For *K_F_* = 0, the estimation is carried out fully in real-time; however, the estimates lack robustness to the fluctuations of the outputs of the attention marker block. The backward-lag *K_B_* incorporates the information before *k*\* in order to make the estimates more reliable, and controls the computational cost of the state-space model for fixed values of *K_F_*. Throughout the rest of the paper, we use the expression *real-time* for referring to algorithms that operate with a fixed forward-lag of *K_F_*. We will discuss specific choices of *K_F_* and *K_B_* and their implications in Section 3.

**Figure 2:**
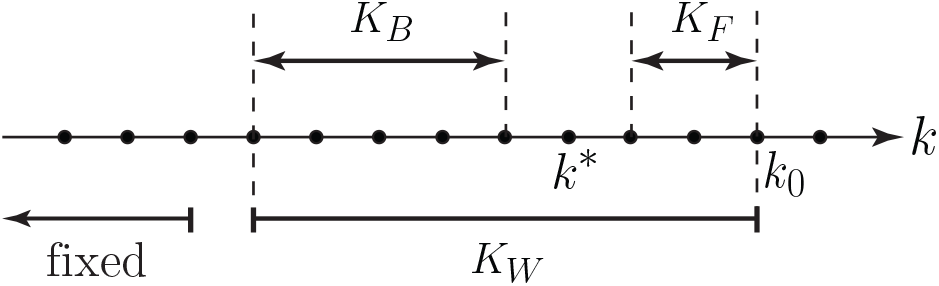
The parameters involved in state-space fixed-lag smoothing.

Suppose we are in a window of length *K_W_* where the instances are indexed by *k* = 1, 2,…, *K_W_*. Inspired by [16], we assume a linear state-space model on the logit-probability of attending to speaker 1. We define the binary random variable *n_k_* = 1 when speaker 1 is attended and *n_k_* = 2 when speaker 2 is attended, at instance *k*. The goal is to obtain estimates of *p_k_* := P (*n_k_* = 1) together with its confidence intervals for 1 ≤ *k* ≤ *K_W_*. The state dynamics are given by:

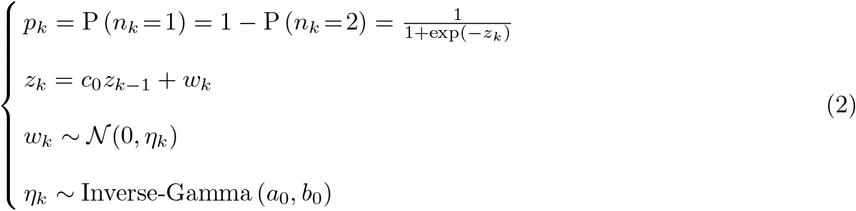

The dynamics of the main latent variable *z_k_* are controlled by its transition scale *c*_0_ and state variance *η_k_*. The hyperparameter 0 ≤ *c*_0_ ≤ 1 ensures the stability of the updates for *z_k_*. The state variance *η_k_* is modeled using an Inverse-Gamma conjugate prior with hyper-parameters *a*_0_ and *b*_0_. The log-prior of the Inverse-Gamma density takes the form 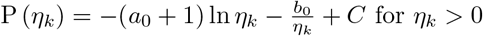, where *C* is a normalization constant. By choosing *a*_0_ greater and sufficiently close to 2, the variance of the Inverse-Gamma distribution takes large values and therefore can serve as a non-informative conjugate prior. Considering the fact that we do not expect the attentional state to have high fluctuations within a small window of time, we can further tune the hyperparameters *a*_0_ and *b*_0_ for the prior to promote smaller values of *η_k_*’s. This way, we can avoid large consecutive fluctuations of the *z_k_*’s, and consequently the *p_k_*’s.

Next, we develop an observation model relating the state dynamics of Eq. (2) to the observations 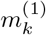 and 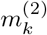 for *k* =1, 2,…, *K_W_*. To this end, we use the latent variable *n_k_* as the link between the states and observations:

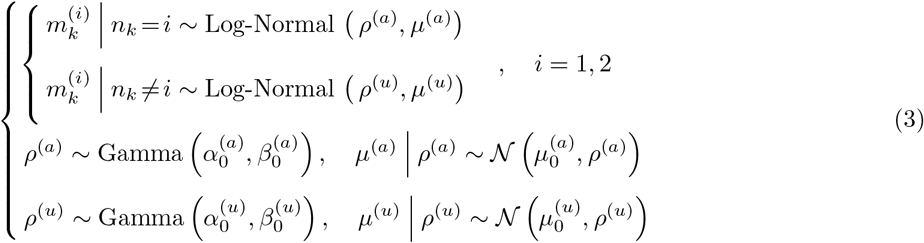

When speaker *i* = 1, 2 is attended to, we use a Log-Normal distribution on 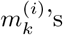, with log-prior given by 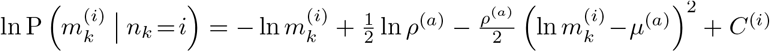, where 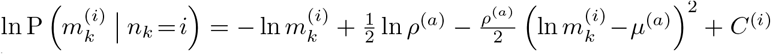, and *C*^(*i*)^ is a normalization constant, for *i* = 1, 2, and *k* = 1, 2,…, *K_W_*. Similarly, when speaker *i* =1, 2 is *not* attended to, we use a Log-Normal prior on 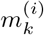 with parameters *ρ*^(*u*)^ and *μ*^(*u*)^. As mentioned before, choosing an appropriate attention marker results in a statistical separation between 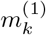 and 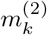, if only one speaker is attended. The Log-Normal distribution is a distribution on 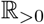 which lets us capture this concentration in the values of 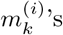. In contrast to [16], this distribution also leads to closed form update rules, which significantly reduces computational costs. We have also imposed conjugate priors on the joint distribution of (*ρ, μ*)’s, which factorizes as lnP(*ρ, μ*) = lnP(*ρ*) + lnP(*μ*|*ρ*). The hyperparameters *α*_0_, *β*_0_, and *μ*_0_ serve to tune the attended and the unattended Log-Normal distributions to create separation between the attended and unattended cases. These hyperparameters can be determined based on the mean and variance information of 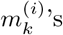 in a supervised manner, where the attended speaker is known.

The parameters of the state-space model are therefore 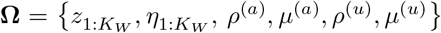, which have to be inferred from 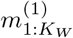 and 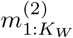. As mentioned before, our goal in the fixed-lag smoothing approach is to estimate *z_k*_* and *η_k*_* in each window, where *k*\* = *K_W_* – *K_F_*. However, in order to do so in our model, we perform the inference step over all the parameters in **Ω** and output the estimates of {*z_k*_, η_k*_*} ∈ **Ω**. The estimated **Ω** would then serve as the initialization for parameter estimation in the next window. The parameters in **Ω** can be inferred through two nested EM algorithms as in [16]. In Section 2 of the Supplementary Material, we have given a detailed derivation of the EM framework and update rules in the real-time setting, as well as solutions to further reduce the computational costs thereof.

*Remark* 4. The state-space models given in Eqs. 2 and 3 have two major differences with the one used in [16]. First, in [16], the distribution over the correlative measure for the *unattended* speaker is assumed to be uniform. However, this assumption may not hold for other attention markers in general. For instance, the M100 magnitude of the TRF estimated from MEG data is a positive random variable, which is concentrated on higher values for the attended speaker compared to the unattended speaker. In order to address this issue, we consider a parametric distribution in Eq. (3) over the attention marker corresponding to the unattended speaker and infer its parameters from the data. If this distribution is indeed uniform and non-informative, the variance of the unattended distribution, which is estimated from the data, would be large enough to capture the flatness of the distribution. Second, the parametrization of the observations using Log-Normal densities and their corresponding priors factorized using Gamma and Gaussian priors, admits fast and closed-form update equations in the real-time setting. As we have shown in Section 2 of the Supplementary Material, these models also have the advantage of incorporating low-complexity updates by simplifying the EM procedure. In addition, the Log-Normal distribution as a generic unimodal distribution allows us to model a larger class of attention markers.

### 2.4. EEG Recording and Experiment Specifications

64-channel EEG was recorded using the actiCHamp system (Brain Vision LLC, Morrisville, NC, US) and active EEG electrodes with Cz channel being the reference. The data was digitized at a 10 KHz sampling frequency. Insert earphones ER-2 (Etymotic Research Inc., Elk Grove Village, IL, US) were used to deliver sound to the subjects while sitting in a sound-attenuated booth. The earphones were driven by the clinical audiometer Piano (Inventis SRL, Padova, Italy), and the volume was adjusted for every subject’s right and left ears separately until the loudness in both ears was matched at a comfortably loud listening level. Three normal-hearing adults participated in the study. The mean age of subjects was 49.5 years with the standard deviation of 7.18 years. The study included a constant-attention experiment, where the subjects were asked to sit in front of a computer screen and restrict motion while any audio was playing. The data used in this paper corresponds to 3 subjects, 24 trials each.

The stimulus set contained eight story segments, each approximately ten minutes long. Four segments were narrated by male speaker 1 (M1) and the other four by male speaker 2 (M2). The stimuli were presented to the subjects in a dichotic fashion, where various stories read by M1 were played in the left ear, while stories read by M2 were played in the right ear for all the subjects. Each subject listened to twenty four trials of the dichotic stimulus. Each trial had a duration of approximately one minute, and for each subject, no storyline was repeated in more than one trial. During each trial, the participants were instructed to look at an arrow at the center of the screen, which determined whether to attend to the right-ear story or to the left one. The arrow remained fixed for the duration of each trial, making it a constant-attention experiment. At the end of each trial, two multiple choice semantic questions about the attended story were displayed on the screen to keep the subjects alert. The responses of the subjects as well as their reaction time were recorded as a behavioral measure of the subjects’ level of attention, and above eighty percent of the questions were answered correctly by each subject. Breaks and snacks were given between stories if requested. All the audio recordings, corresponding questions, and transcripts were obtained from a collection of stories recorded at Hafter Auditory Perception Lab at UC Berkeley.

### 2.5 MEG Recording and Experiment Specifications

MEG signals were recorded with a sampling rate of 1 KHz using a 160-channel whole-head system (Kanazawa Institute of Technology, Kanazawa, Japan) in a dimly lit magnetically shielded room (Yokogawa Electric Corporation). Detection coils were arranged in a uniform array on a helmet-shaped surface on the bottom of the dewar with 25 mm between the centers of two adjacent 15.5 mm diameter coils. Also, sensors are set as first-order axial gradiometers with a baseline of 50 mm, resulting in field sensitivities of 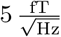 or better in the white noise region.

The two speech signals had approximately 65 dB SPL and were presented using the software package Presentation (Neurobehavioral Systems Inc., Berkeley, CA, US). The stimuli were delivered to the subjects’ *Õ* ears with 50Ω sound tubing (E-A-RTONE 3A; Etymotic Research), attached to E-A-RLINK foam plugs inserted into the ear canal. Also, the whole acoustic delivery system was equalized to give an approximately flat transfer function from 40 Hz to 3000 Hz. A 200 Hz low-pass filter and a notch filter at 60 Hz were applied to the magnetic signal in an online fashion for noise removal. Three of the 160 channels were magnetometers separated from the others and used as reference channels. Finally, to quantify the head movement, five electromagnetic coils were used to measure each subject’s head position inside the MEG machine once before and once after the experiment.

Nine normal-hearing, right-handed young adults (ages between 20 and 31) participated in this study. The study includes two sets of experiments: the constant-attention experiment and the attention-switch experiment, in both of which six subjects participated. Three subjects took part in both of the experiments. The experimental procedure were approved by the University of Maryland Institutional Review Board (IRB), and written informed consent was obtained from each subject before the experiment.

The stimuli included four non-overlapping segments from the book *A Child’s History of England* by Charles Dickens. Two of the segments were narrated by a man and the other two by a woman. Three different mixtures, each 60 s long, were generated and used in the experiments to prevent reduction in the attentional focus of the subjects. Each mixture included a segment narrated by the male speaker and one narrated the the female speaker. In all trials, the stimuli were delivered diotically to both ears using tube phones inserted into the ear canals at a roughly 65 dB SPL, as mentioned. The constant-attention experiment consisted of two conditions: 1) attending to the male speaker in the first mixture, 2) attending to the female speaker in the second mixture. In the attention-switch experiment, subjects were instructed to focus on the female speaker in the first 28s of the trial, switch their attention to the male speaker after hearing a 2s pause (28th to 30th seconds), and maintain their focus on the latter speaker through the end of the trial. Each mixture was repeated three times in the experiments, resulting in six trials per speaker for the constant-attention experiment and three trials per speaker for the attention-switch experiment. After the presentation of each mixture, subjects answered comprehensive questions related to the segment they were instructed to focused on, as a way to keep them motivated on attending to the target speaker. Eighty percent of the questions were answered correctly on average. Furthermore, a pilot study for each of the nine participating subjects was performed prior to the main experiments. In this study, the subjects listened to a single speech stream, first segment in the stimuli set narrated by the male speaker, for three trials each 60 s long. The MEG recordings in the pilot study were used to calculate the subject-specific linear combination of MEG channels which forms the auditory component of the response, as will be explained next. Note that for each subject, all the recordings were performed in a single session resulting in a minimal change of the subject’s head position with respect to the MEG sensors.

## 3. Results

In this section, we apply our real-time attention decoding framework to synthetic data as well as M/EEG recordings. Subsection 3.1 includes the simulation results, and subsections 3.2 and 3.3 demonstrate the results for the analysis of EEG and MEG recordings, respectively.

### 3.1 Simulations

In order to validate our proposed framework, we perform two sets of simulations. The first simulation pertains to our EEG analysis and employs a decoding model, which we will describe below in full detail. The second simulation for our MEG analysis using an encoding model, is deferred to the Supplementary Material Section 3, in the interest of space.

#### 3.1.1 Simulation Settings

In order to simulate EEG data under a dual-speaker condition, we use the following generative model:

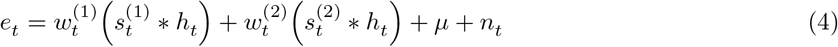

where 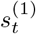 and 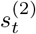 are respectively the speech envelopes of speakers 1 and 2 at time *t*; the output *e_t_* is the neural response, which denotes an auditory component of the EEG recordings or the measured EEG response at a given channel at time *t* for *t* = 1, 2,…, *T*. Motivated by the analysis of LTI systems, *h_t_* can be considered as the impulse response of the neural process resulting in *e_t_*, and * represents the convolution operator; the scalar *μ* is an unknown constant mean, and *n_t_* denotes a zero-mean i.i.d Gaussian noise. The weight functions 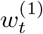 and 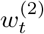 are signals modulated by the attentional state which determine the contributions of speakers 1 and 2 to *e_t_*, respectively. In order to simulate the attention modulation effect, we assume that when speaker 1 (resp. 2) is attended to at time *t*, we have 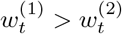 (resp. 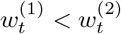).

We have chosen two 60 s-long speech segments from those used in the MEG experiment (See section 2.5) and calculated 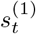 and 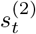 as their envelopes for a sampling rate of *f_s_* = 200 Hz. Also, we have set *μ* = 0.02 and 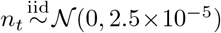 in Eq. (4). Fig. 3-A shows the location and amplitude of the lag components in the impulse response, which is then smoothed using a Gaussian kernel with standard deviation of 10 ms to result in the final impulse response *h_t_*, shown in Fig. 3-B. The significant components of *h_t_* are chosen at 50 ms and 100 ms lags, with few smaller components at higher latencies [16]. The weight signals 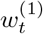 and 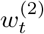 in Eq. (4) are chosen to favor speaker 1 in the [0s, 30s) interval and speaker 2 in the (30s, 60s] interval, with the transition happening within a 3 s interval around the 30 s mark.

**Figure 3:**
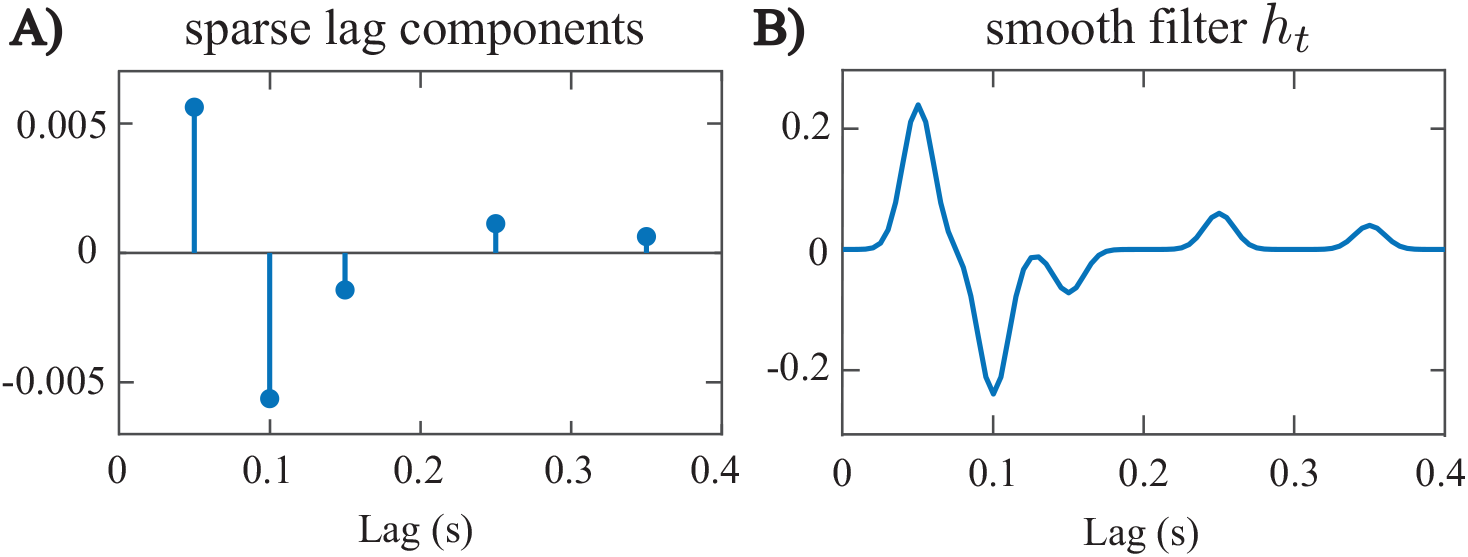
Impulse response *h_t_* used in Eq. (4). A) sparse lag components, B) the smooth impulse response.

#### 3.1.2 Parameter Selection

We aim at estimating decoders in this simulation, which linearly map **e**_*t*_ and its lags to 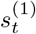 and 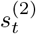. To estimate the decoders, we have considered consecutive non-overlapping windows of length 0.25 s resulting in *K* = 240 windows of length *W* = 50 samples. Also, we have chosen *γ* = 0.001 through cross-validation and λ = 0.95 in estimating the decoding coefficients, which results in an *effective* data length of 5s for decoder estimation. The forward lags of the neural response have been limited to a 0.4 s window, i.e., *L_d_* = 80 samples. Given that the decoder corresponds to the inverse of a smooth kernel *h_t_*, it may not have the same smoothness properties of *h_t_*. Hence, we do not employ a smooth basis for decoder estimation. We have used the FASTA package [29] with Nesterov’s acceleration method to implement the forward-backward splitting algorithm for encoder/decoder estimation. As for the state-space model estimators, we have considered 20 (inner and outer) EM iterations for the batch-mode estimates that use the entire data, while for the real-time estimates, we use 1 inner EM iteration and 20 outer EM iterations (See Section 2 of the Supplementary Material for more details).

There are three criteria for choosing the fixed-lag smoothing parameters: First, how close to the true real-time analysis the system operates is determined by *K_F_*. Second, the computational cost of the system is determined by *K_W_*. Third, how close the output of the system is to that of batch-mode processing is determined by both *K_F_* and *K_W_*. These three criteria form a tradeoff in tuning the parameters *K_W_* and *K_F_*. Specific choices of these parameters are given in the next subsection.

For tuning the hyperparameters of the priors on the attended and unattended distributions, we have used a separate 15 s sample trial generated from the same simulation model in Eq. (4) for each of the three cases. The parameters 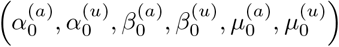 have been chosen by fitting the Log-Normal distributions to the attention marker outputs from the sample trials in a supervised manner (with known attentional state). The variance of the Gamma priors 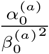 and 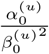 have been chosen large enough such that the priors are non-informative. This step can be thought of as the initialization of the algorithms prior to data analysis. For the Inverse-Gamma prior on the state-space variances, we have chosen *a*_0_ = 2.008 and *b*_0_ = 0.2016, resulting in a mean of 0.2 and a variance of 5. This prior favors small values of *η_k_*’s to ensure that the state estimates are immune to large fluctuations of the attention markers, while the large variance (compared to the mean) results in a non-informative prior.

#### 3.1.3 Estimation Results

Fig. 4 shows the results of our estimation framework for a correlation-based attention marker. Row A in Fig. 4 shows three cases considered for modulating the weights 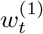 and 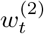, where the weights are contaminated with Gaussian noise 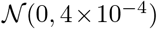. Cases 1, 2, and 3 exhibit increasing levels of difficulty in discriminating the contributions of the two speakers to the neural response. Rows B and C in Fig. 4 respectively show the decoder estimates for speakers 1 and 2. As expected, the significant components of the decoders around 50 ms, 100 ms, and 150 ms lags, are modulated by the attentional state, and the modulation effect weakens as we move from Case 1 to 3. In Case 1, these components are less significant overall for the decoder estimates of speaker 2 in the [0 s, 30 s] time interval and become larger as the attention switches to speaker 2 during the rest of the trial (red boxes in row C of Case 1). On the other hand, in Case 3, the magnitude of the said components do not change notably across the 30 s mark.

**Figure 4:**
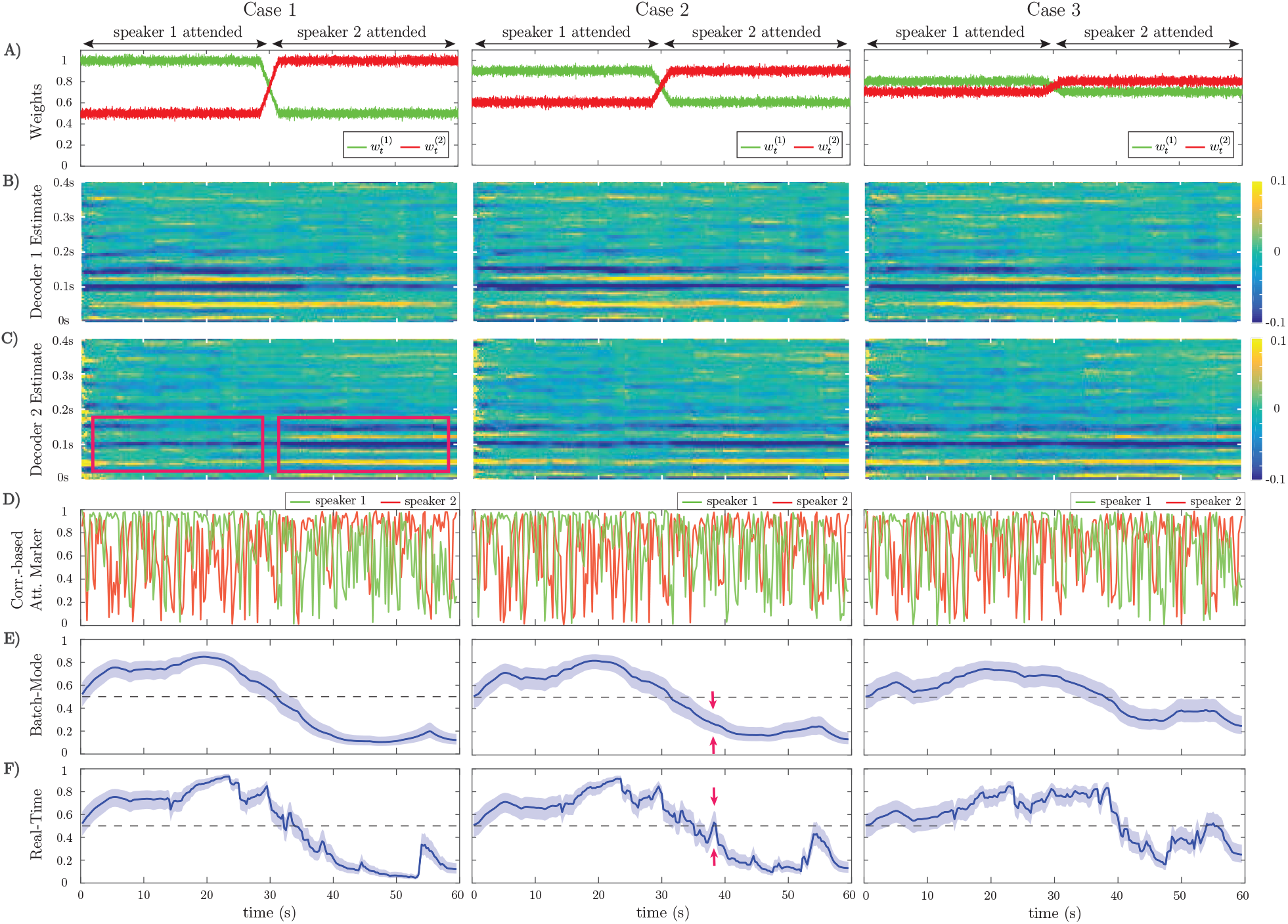
Estimation results of application to simulated EEG data for the correlation-based attention marker: A) Input weights 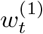 and 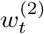 in Eq. (4), which determine the relative effect of the two speeches on the neural response. Based on our generative model, the attention is on speaker 1 for the first half of each trial and on speaker 2 for the second half. Case 1 corresponds to a scenario where the effects of the attended and unattended speeches in the neural response are well-separated. This separation decreases as we move from Case 1 to Case 3. B) Estimated decoder for speaker 1. C) Estimated decoder for speaker 2. In Case 1, the significant components of the estimated decoders near the 50 ms, 100 ms, and 150 ms lags are notably modulated by the attentional state as highlighted by the red boxes. This effect weakens in Case 2 and visually disappears in Case 3. D) Output of the correlation-based attention marker for each speaker. E) Output of the batch-mode state-space estimator for the correlation-based attention marker as the estimated probability of attending to speaker 1. F) Output of the real-time state-space estimator, i.e., fixed-lag smoother, for the correlation-based attention marker as the estimated probability of attending to speaker 1. The real-time estimator is not as robust as the batch-mode estimator to the stochastic fluctuations of the attention marker in row D and is more prone to misclassifications. The red arrows in rows E and F of Case 2 show that the batch-mode estimator correctly classifies the instance as attending to speaker 2, while the real-time estimator is unable to determine the attentional state.

We have considered two different attention markers for this simulation. Row D in Fig. 4 displays the output of a correlation-based attention marker for speakers 1 and 2, which is calculated as 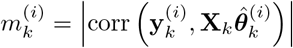 for *i* = 1, 2 and *k* = 1,2, …, *K*. As discussed in subsection 2.2, this attention marker is a measure of how well a decoder can reconstruct its target envelope. As observed in row D of Fig. 4, the attention marker is a highly variable surrogate of the attentional state at each instance, i.e., *on average* the attention marker output for speaker 1 is higher then that of speaker 2 in the [0 s, 30 s) interval and vice versa in the (30 s, 60 s] interval. The reliability of the attention marker significantly degrades going from Case 1 to 3. This highlights the need for state-space modeling and estimation in order to optimally exploit the attention marker.

Rows E and F in Fig. 4 respectively show the batch-mode and real-time estimates of the attentional state probabilities *p_k_* = P (*n_k_* = 1) for *k* = 1,…, *K*, for the correlation-based attention marker, where colored halls indicate 90% confidence intervals. Row F in Fig. 4 corresponds to the fixed-lag smoother, using a window of length 15s (*K_W_* = ⌊15*f_s_*/*W*⌋), and a forward-lag of 1.5s (*K_F_* = ⌊1.5*f_s_*/*W*⌋). We refer to this estimator as the real-time estimator henceforth. Note that by accounting for the forward-lag in the decoder (*L_d_*), the overall delay in estimating the attentional state is 1.9 s. Recall that in batch-mode processing, all of the attention marker outputs across the trial are available the state-space estimator, as opposed to the fixed-lag estimator which has access to a limited number of the attention markers. Therefore, the output of the batch-mode estimator (Row E) is a more robust measure of the instantaneous attentional state as compared to the real-time estimator (Row F), since it is less sensitive to the stochastic fluctuations of the attention markers in row D. For example, in the instance marked by the red arrows in rows E and F of Case 2 in Fig. 4, the batch-mode estimator classifies the instance correctly as attending to speaker 2, while the real-time estimator cannot make an informed decision since *p_k_* = 0.5 falls within the 90% confidence interval of the estimate at this instance. However, the real-time estimator exhibits performance closely matching that of the batch-mode estimator for most instances, while operating in real-time with limited data access and significantly lower computational complexity.

Row A in Fig. 5 exhibits the output of another attention marker computed as the ℓ_1_-norm of the decoder given by 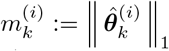 for *i* = 1, 2 and *k* = 1, 2,…, *K*, where the first element of 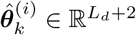 (the intercept parameter) is discarded in computing the ℓ_1_-norm. This attention marker captures the effect of the significant peaks in the decoder. The rationale behind using the ℓ_1_-norm based attention marker is the following: in the extreme case that the neural response is solely driven by the attended speech, we expect the unattended decoder coefficients to be small in magnitude and randomly distributed across the time lags. The attended decoder, however, is expected to have a sparse set of informative and significant components corresponding to the specific latencies involved in auditory processing. Thus, the ℓ_1_ norm serves to distinguish between these two cases. Rows B and C in Fig. 5 show the batch-mode and real-time estimates of the attentional state probabilities for the ℓ_1_-norm attention marker, respectively, where colored halls indicate 90% confidence intervals. Consistent with the results of the correlation-based attention marker (Rows E and F in Fig. 4), the real-time estimator exhibits performance close to that of the batch-mode estimator. Comparing Figs. 4 and 5 reveals the dependence of the attentional state estimation performance on the choice of the attention marker: while the correlation-based attention marker is more widely used, the ℓ_1_-based attention marker provides smoother estimates of the attention probabilities, and can be used as a more robust alternative to the correlation-based attention marker.

**Figure 5:**
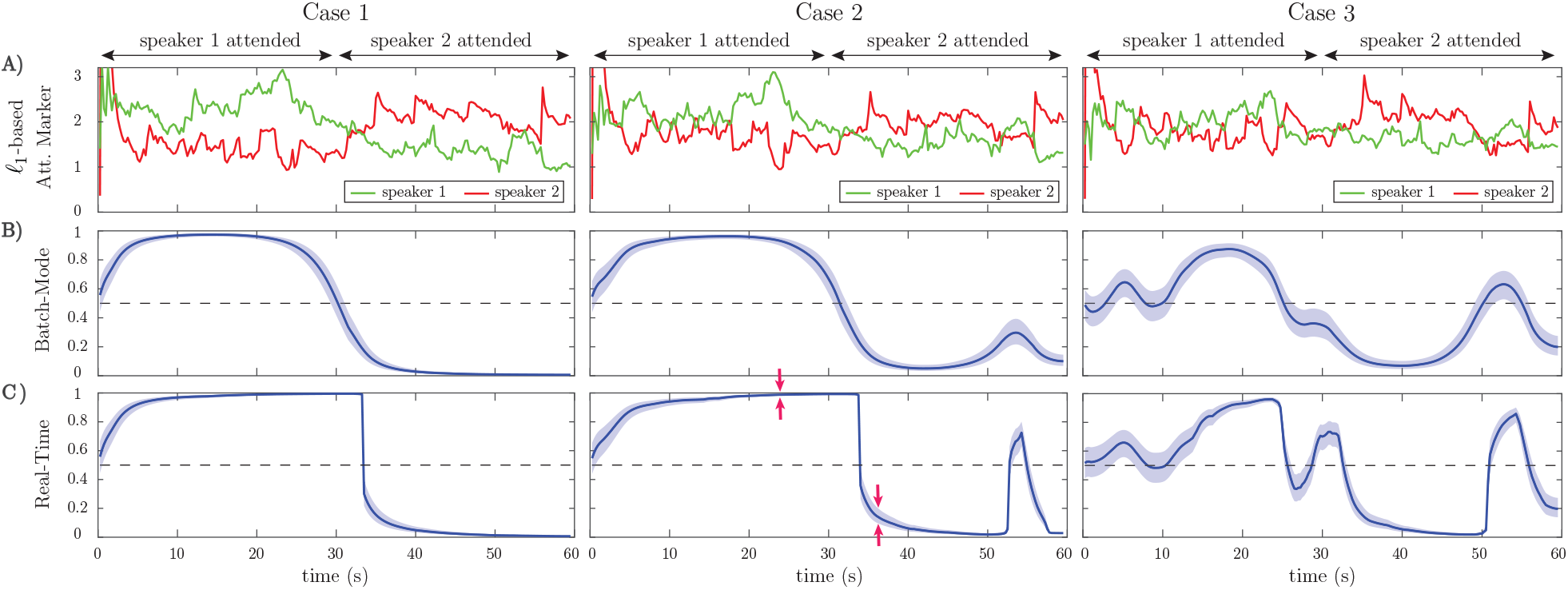
Estimation results of application to simulated EEG data for the ℓ_1_-based attention marker: A) Output of the ℓ_1_-based attention marker for each speaker, corresponding to the three cases in Figure 4. B) Output of the batch-mode state-space estimator for the ℓ_1_-based attention marker as the estimated probability of attending to speaker 1. C) Output of the real-time state-space estimator for the ℓ_1_-based attention marker as the estimated probability of attending to speaker 1. Similar to the preceding correlation-based attention marker, the classification performance degrades when moving from Case 1 (strong attention modulation) to Case 3 (weak attention modulation).

#### 3.1.4 Discussion and Further Analysis

Going from Case 1 to Case 3 in Fig. 4 and Fig. 5, we observe that the performance of all estimators degrades, causing a drop in the classification accuracy and confidence. This performance degradation is due to the declining power of the attention markers in separating the contributions of the attended and unattended speakers. However, comparing the outputs of the real-time and batch-mode estimators with their corresponding attention marker outputs in row D of Fig. 4 and row A of Fig. 5, highlights the role of the state-space model in suppressing the stochastic fluctuations of the attention markers and thereby providing a robust and smooth measure of the attentional state.

It is noteworthy that all the estimators exhibit a systematic delay in detecting the deflection point at 30 s, even for the well-separated Case 1 and batch-mode estimation. This delay is due to two main factors: first, the transition period of 3 s in the design of the weight signals contributes to this delay. Second, although the forgetting factor mechanism used in estimating the decoder coefficients results in more stable estimates, it causes an extra delay to the overall performance of the estimator.

Comparing the batch-mode and the real-time estimators in Fig. 4 and Fig. 5, we observe that the real-time estimators closely follow the output of the batch-mode estimators, while having access to data in an online fashion. A significant deviation between the batch-mode and real-time performance is observed in rows B and C (Cases 1 and 2) of Fig. 5 in the form of sharp drops in the real-time estimates of the attentional state probability. Given that the real-time estimator has only access to the attention marker within *K_F_* samples in the future, the confidence intervals significantly narrow down within the first half of the trial, as all the past and near-future observations are consistent with attention to speaker 1. However, shortly after the 30 s mark the estimator detects the change and the confidence bounds widen accordingly (see red arrows in row C of Case 2 in Fig. 5).

In order to further quantify the performance gap between the batch-mode and real-time estimators, we define their relative Mean Squared Error (MSE) as:

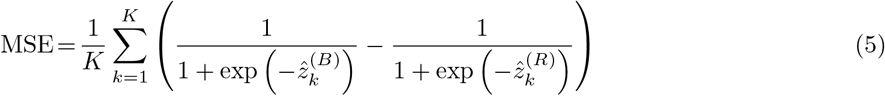

where 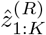 and 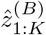 denote the real-time and batch-mode state estimates over a given trial, respectively. We have considered the logistic transformation of 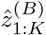 and 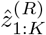, which gives the probability of attending to speaker 1.

Figure 6 shows the effect of varying the forward-lag *K_F_* from 0s (i.e., fully real-time) to 5s with 0.5s increments for the two attention markers in Case 2 of Fig. 4 and Fig. 5, as an example. All of the other parameters in the simulation have been fixed as before. The left panels in Fig. 6 show the MSE for different values of *K_F_* in the real-time setting. As expected, for both attention markers, the MSE decreases as the forward-lag increases. The right panels in Fig. 6 display the incremental MSE defined as the change in MSE when *K_F_* is increased by 0.5s, starting from *K_F_* = 0s. Notice that even a 0.5s forward-lag significantly decreases the MSE from K_F_ = 0s. The subsequent improvements of the MSE diminish as *K_F_* is increased further. Our choice of *K_F_* = 1.5 s in the foregoing analysis was made to maintain a reasonable tradeoff between the MSE improvement and the delay in real-time operation.

**Figure 6:**
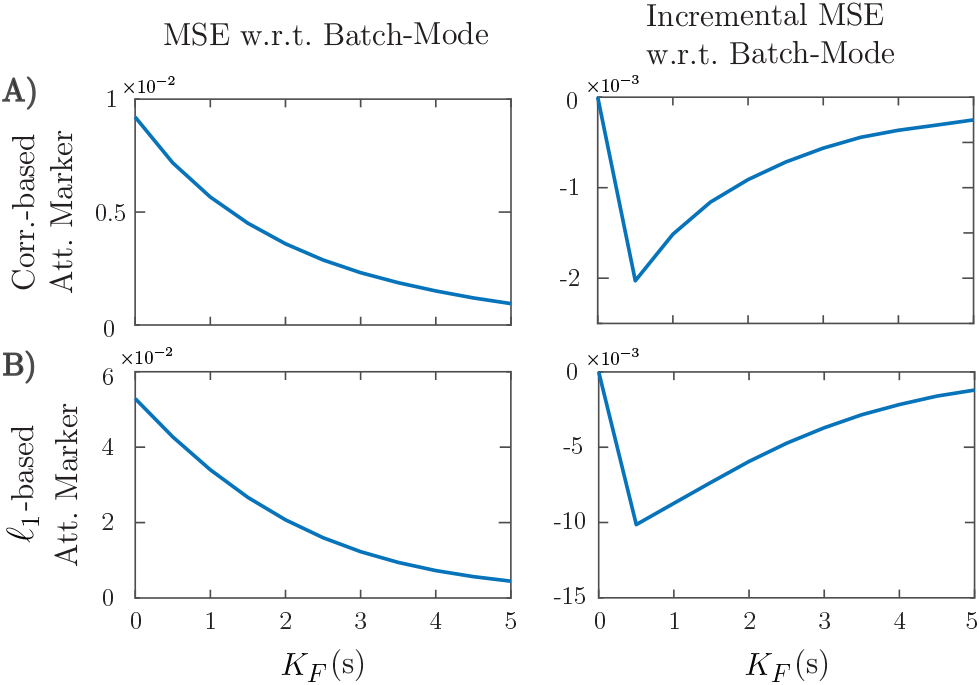
Effect of the forward-lag *K_F_* on the MSE for the two attention markers in case 2 of Fig. 4 and Fig. 5. A) Correlation-based attention marker, B) ℓ_1_-based attention marker. As the forward-lag increases, the MSE decreases, and the output of the real-time estimator becomes more similar to that of the batch-mode. This results in more robustness for the real-time estimator at the expense of more delay in decoding the attentional state. The right panels show that the incremental improvement to the MSE decreases as *K_F_* increases.

Finally, Fig. 7 shows the estimated attention probabilities and their 90% confidence intervals for the correlation-based attention marker in Case 2 of Fig. 4, as an example. The three curves correspond to the extreme values of *K_F_* in Fig. 6 given by *K_F_* = 0s (blue) and *K_F_* = 5s (red), and the batch-mode estimate (green). All the other parameters have been fixed as explained before. The fixed-lag smoothing approach with *K_F_* = 5s is as robust as the batch-mode estimate. The fully real-time estimate with *K_F_* = 0s follows the same trend as the other two. However, it is susceptible to the stochastic fluctuations of attention marker, which may lead to misclassifications (see the red arrows in Fig. 7).

**Figure 7:**
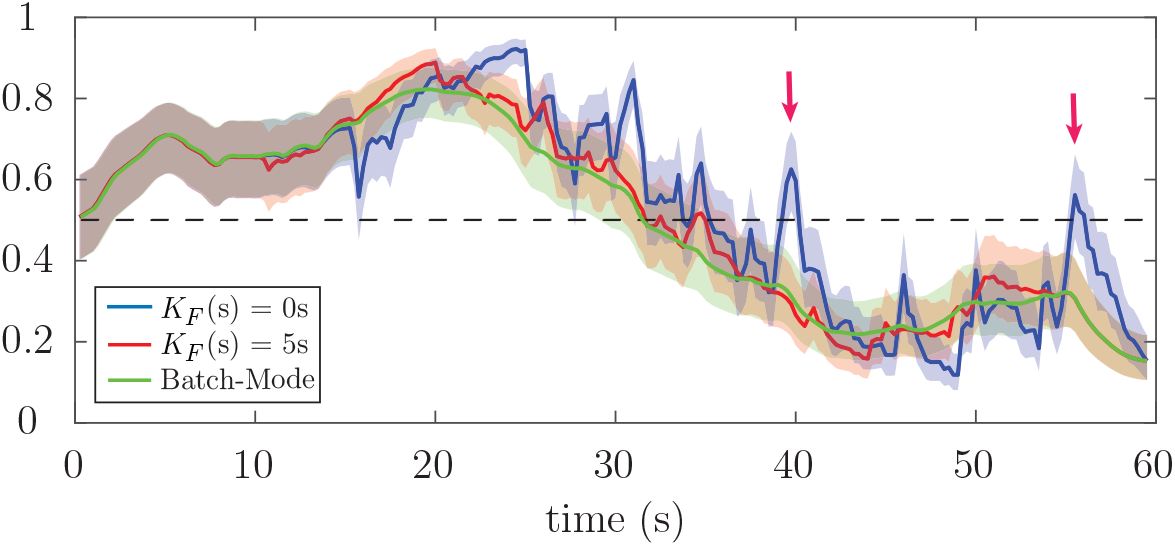
Estimated attention probabilities together with their 90% confidence intervals for the correlation-based attention marker in Case 2 of Fig. 4. The blue, red and green curves correspond to *K_F_* = 0s, *K_F_* = 5s, and batch-mode estimation, respectively. The estimator for *K_F_* = 5s is nearly as robust as the batch-mode. However, the fully real-time estimator with *K_F_* = 0s is sensitive to the stochastic fluctuations of the attention markers, which results in the misclassification of the attentional state at the instances marked by red arrows.

### 3.2. Application to EEG

In this subsection, we apply our real-time attention decoding framework to EEG recordings in a dual-speaker environment. Details of the experimental procedures are given in Section 2.4.

#### 3.2.1 Preprocessing and Parameter Selection

Both the EEG data and the speech envelopes were downsampled to *f_s_* = 64 Hz using an anti-aliasing filter. As the trials had variable lengths, we have considered the first 53 s of each trial for analysis. We have considered consecutive windows of length 0.25 s for decoder estimation, resulting in *W* = 16 samples per window and *K* = 212 instances for each trial. Also, we have considered lags up to 0.25 s for decoder estimation, i.e., *L_d_* = 16. The latter is motivated by the results of [10] suggesting that the most relevant decoder components are within the first 0.25 s lags. Prior studies have argued that the effects of auditory attention and speech perception are strongest in the frontal and close-to-ear EEG electrodes [11, 31, 32, 33]. We have only considered 28 EEG channels in the decoder estimation problem, i.e., *C* = 28, including the frontal channels Fz, F1-F8, FCz, FC1-FC6, FT7-FT10, C1-C6, and the T complex channels T7 and T8. According to [12], using only this number of electrodes in the decoding process results in nearly the same classification performance as in the case of using all the electrodes. Note that for our real-time setting, a channel selection step can considerably decrease the computational cost and the dimensionality of the decoder estimation step, given that a vector of size 1 + *C*(*L_d_* + 1) needs to be updated within each 0.25s window.

We have determined the regularization coefficient *γ*=0.4 via cross-validation and the forgetting factor λ = 0.975, which results in an *effective* data length of 10 s in the estimation of the decoder and is long enough for stable estimation of the decoding coefficients. It is worth noting that small values of λ, and hence small effective data lengths, may result in an under-determined inverse problem, since the dimension of the decoder is given by 1 + *C*(*L_d_* +1). Finally, in the FASTA package, we have used a tolerance of 0.01 together with Nesterov’s accelerated gradient descent method to ensure that the processing can be done in an online fashion.

In studies involving correlation-based measures, such as [10, 16], the convention is to train attended and unattended decoders/encoders using multiple trials and then use them to calculate the correlation measures over the test trials. The correlation-based attention marker, however, did not produce a statistically significant segregation of the attended and the unattended speakers in our analysis. This discrepancy seems to stem from the fact that the estimated encoders/decoders and the resulting correlations in the aforementioned studies are more informative and robust due to the use of batch-more analysis with multiple trials, as compared to our real-time framework. The ℓ_1_-based attention marker, however, resulted in a meaningful statistical separation between the attended and the unattended speakers. Therefore, in what follows, we present our EEG analysis results using the ℓ_1_-based attention marker.

The parameters of the state-space models have been set similar to those used in simulations, i.e., *K_W_* = ⌊15*f_s_*/*W*⌋, *K_F_* = ⌊1.5*f_s_*/*W*⌋, *a*_0_ = 2.008, *b*_0_ = 0.2016. Considering the 0.25s lag in the decoder model, the total delay in estimating the attentional state for the real-time system is 1.75 s. For estimating the prior distribution parameters for each subject, we use the first 15s of each trial. As mentioned before, considering the 15 s-long sliding window, we can treat the first 15 s of each trial as a tuning step in which the prior parameters are estimated in a supervised manner and the state-space model parameters are initialized with the values estimated using these initial windows. Thus, similar to the simulations, 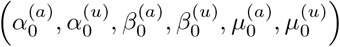 for each subject have been set according to the parameters of the two fitted Log-Normal distributions on the ℓ_1_-norm of the decoders in the first 15 s of the trials, while choosing large variances for the priors to be non-informative.

#### 3.2.2. Estimation Results

Fig. 8 shows the results of applying our proposed framework to EEG data. For graphical convenience, the data have been rearranged so that speaker 1 is always attended. The left, middle and right panels correspond to subjects 1, 2, and 3, respectively. For each subject, three example trials have been displayed in rows A, B, and C. Row A includes trials in which the attention marker clearly separates the attended and unattended speakers, while Row C contains trials in which the attention marker fails to do so. Row B displays trials in which on average the ℓ_1_-norm of the estimated decoder is larger for the attended speaker; however, occasionally, the attention marker fails to capture the attended speaker.

**Figure 8:**
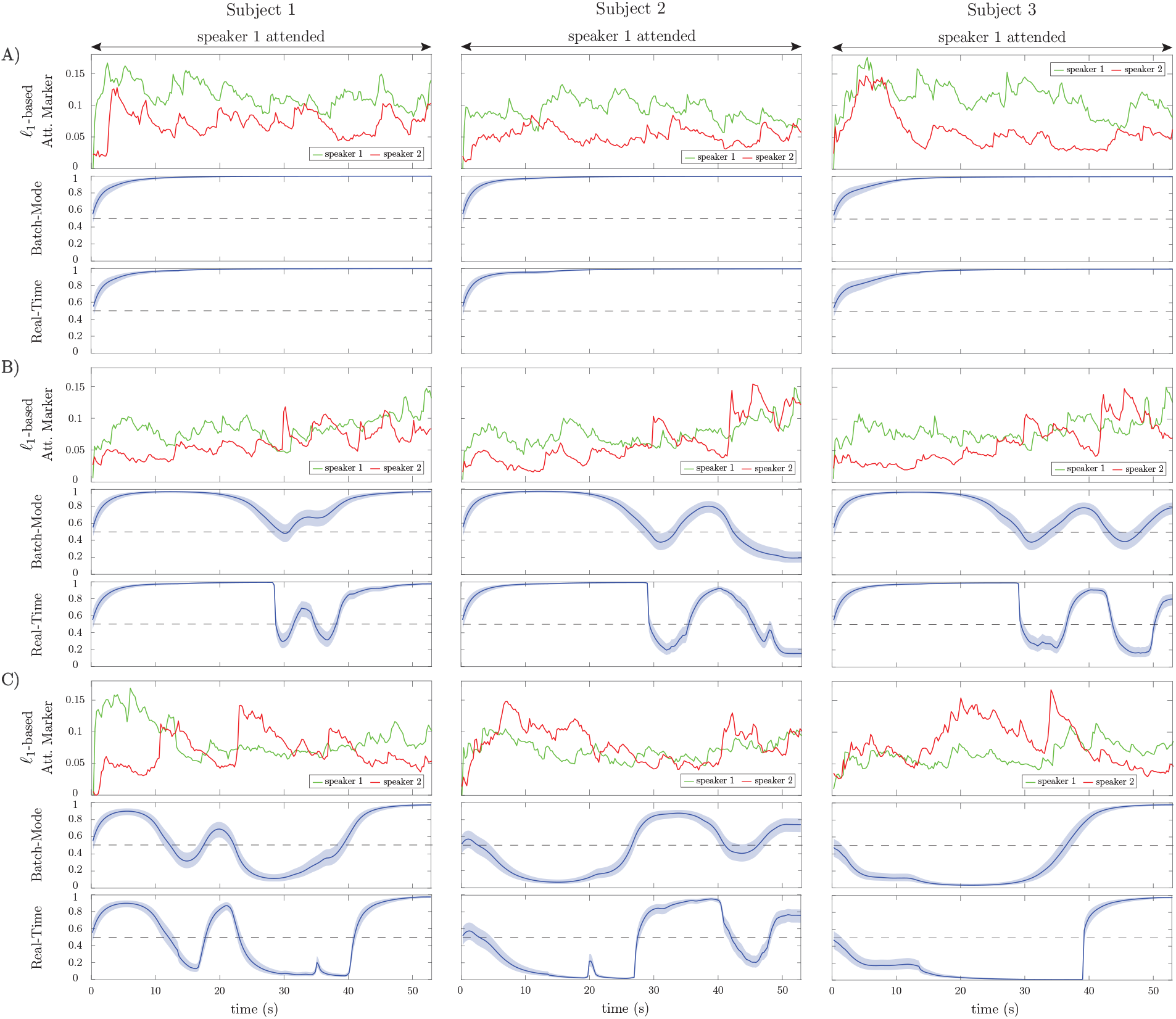
Examples of the ℓ_1_-based attention markers (first panels), batch-mode (second panels), and real-time (third panels) state-space estimation results for nine selected EEG trials. A) Representative trials in which the attention marker reliably separates the attended and unattended speakers. B) Representative trials in which the attention marker separates the attended and unattended speakers on average over the trial. C) Representative trials in which the attention marker either does not separate the two speakers or results in a larger output for the unattended speaker.

Consistent with our simulations, the real-time estimates (third graphs in rows A, B and C) generally follow the output of the batch-mode estimates (second graphs in rows A, B and C). However, the batch-mode estimates yield smoother transitions and larger confidence intervals in general, both of which are due to having access to future observations.

Figure 9 shows the effect of forward-lag *K_F_* on the performance of real-time estimates, similar to that shown in Fig. 6 for the simulations. The forward-lag *K_F_* is increased from 0s to 5s with 0.5 s increments while all the other parameters of the EEG analysis remain the same. The MSE in Fig. 9 has been averaged over all trials for each subject. As we observe in the incremental MSE plot, even a 0.5s lag can significantly decrease the MSE from the case of *K_F_* = 0s (corresponding to the fully real-time setting). Similar to the simulations, we have chosen *K_F_* = 1.5s for the EEG analysis, since the incremental MSE improvements are significant at this lag, and this choice results in a tolerable delay for real-time applications.

**Figure 9:**
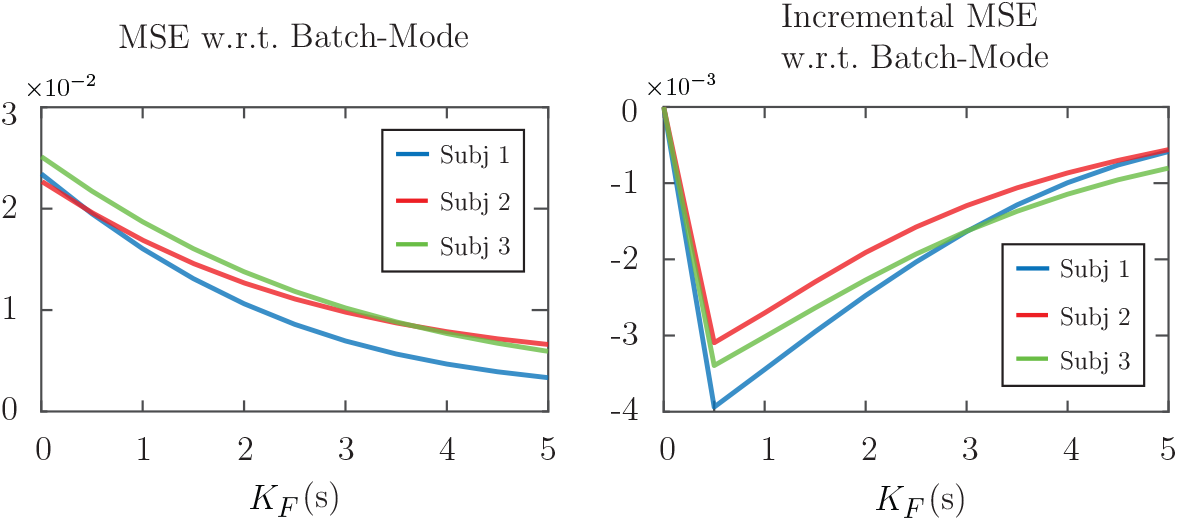
Effect of the forward-lag *K_F_* on MSE in application to real EEG data. The left panel shows the MSE with respect to the batch-mode output averaged over all the trials for each subject. The right panel displays the incremental MSE at each lag, from *K_F_* =0s to *K_F_* = 5s with 0.5 s increments.

Finally, Fig. 10 summarizes the *real-time* classification results of our EEG analysis at the group level.

**Figure 10:**
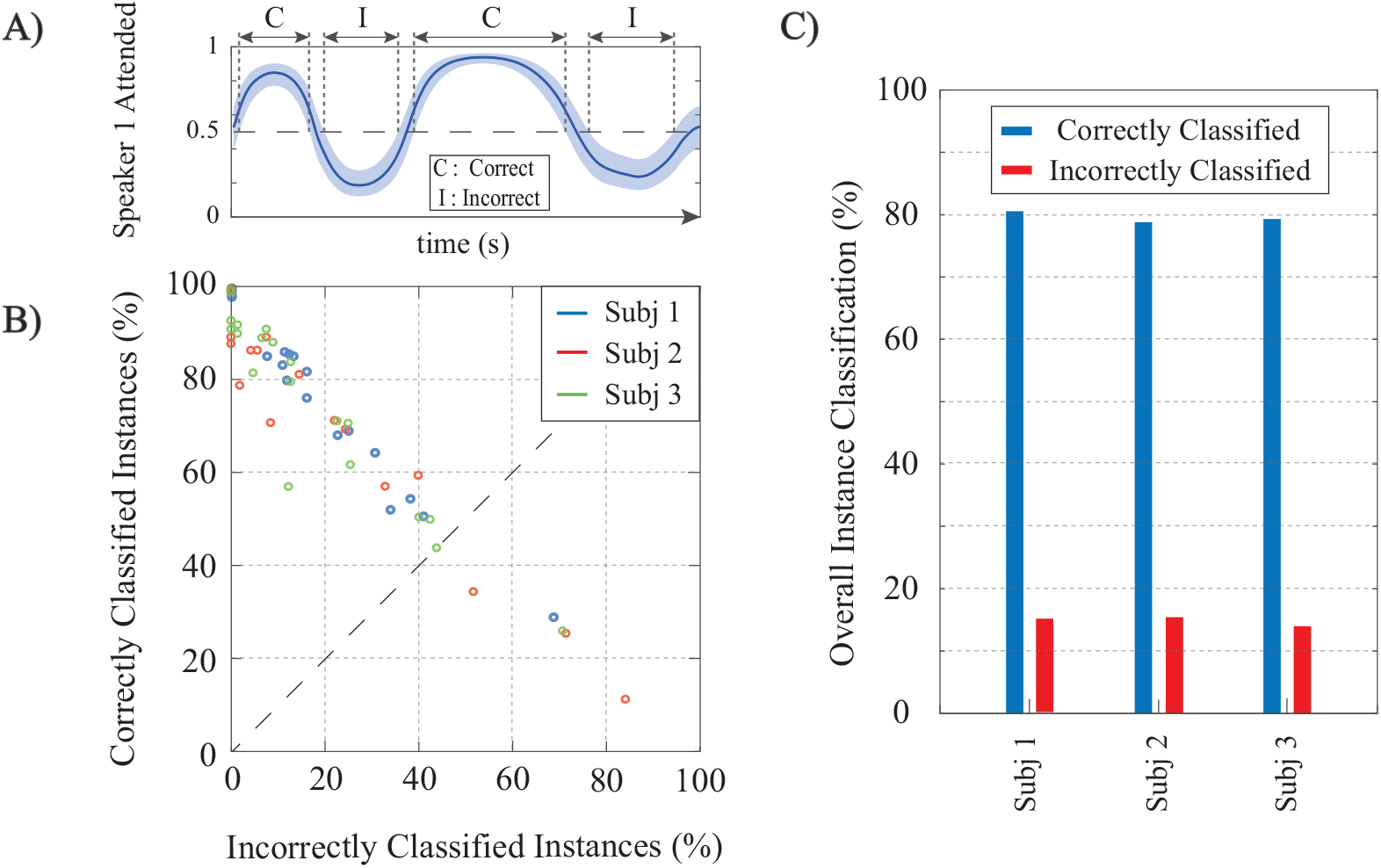
Summary of the real-time classification results in application to real EEG data. A) a generic example of the state-space output for a trial illustrating the classification conventions. B) Classification results per trial for all subjects; each circle corresponds to a trial and the subjects are color-coded. The trials falling below the dashed line have more incorrectly classified instances than correctly classified ones. C) Average classification performance over all trials for the three subjects.

Fig. 10-A shows a cartoon of the estimated attention probabilities for a generic trial in order to illustrate the classification conventions. We define an instance (i.e., *K* consecutive windows of length *W*) to be correctly (incorrectly) classified if the estimated attentional state probability together with its 90% confidence intervals lie above (below) 0.5. If the 90% confidence interval at an instance includes the 0.5 attention probability line, we do not classify it as either correct or incorrect. Figure 10-B displays the correctly classified instances (y-axis) versus those incorrectly classified (x-axis) for each trial. The subjects are color-coded and each circle corresponds to one trial. The average classification results over all trials for each subject are shown in Figure 10-C. In summary, our framework provides ~ 80% average hit rate and ~ 15% average false-alarm per trial per subject. The group-level hit rate and false alarm rate are respectively given by 79.63% and 14.84%.

### 3.3 Application to MEG

In this subsection, we apply our real-time attention decoding framework to MEG recordings of multiple subjects in a dual-speaker environment. The MEG experimental procedures are discussed in Section 2.5.

#### 3.3.1 Preprocessing and Parameter Selection

The recorded MEG responses were band-pass filtered between 1 Hz-8 Hz (delta and theta bands), corresponding to the slow temporal modulations in speech [14, 13], and downsampled to 200 Hz. MEG recordings, like EEG, include both the stimulus-driven response as well as the background neural activity, which is irrelevant to the stimulus. For the encoding model used in our analysis, we need to extract the stimulus-driven portion of the response, namely the auditory component. In [34, 21], a blind source separation algorithm called the Denoising Source Separation (DSS) has been introduced which decomposes the data into temporally uncorrelated components ordered according to their trial-to-trial phase-locking reliability. In doing so, DSS only requires the responses in different trials and not the stimuli. Similar to [17, 16], we only use the first DSS component as the auditory component, since it tends to capture a significant amount of stimulus information and to produce a bilateral stereotypical auditory field pattern.

Since DSS is an *offline* algorithm operating on all the data at once, we cannot readily use it for real-time attention decoding. Instead, we apply DSS to the data from pilot trials from each subject in order to calculate the *subject-specific* linear combination of the MEG channels that compose the first DSS component. We then use these channel weights to extract the MEG auditory responses during the constant-attention and attention-switch experiments in a real-time fashion. Note that the MEG sensors are not fixed with respect to the head position across subjects and are densely distributed in space. Therefore, it is not reasonable to use the same MEG channel weights for all subjects. The pilot trials for each subject can thus serve as a training and tuning step prior to the application of our proposed attention decoding framework.

The MEG auditory component extracted using DSS is used as *E_t_* in our encoding model. Similar to our foregoing EEG analysis, we have considered consecutive windows of length 0.25 s resulting in *W* = 50 samples per window and a total number of *K* = 240 instances, at a sampling frequency of 200 Hz. The TRF length, or the total encoder lag, has been set to 0.4 s resulting in *L_e_* = 80 in order to include the most significant TRF components [13]. The ℓ_1_-regularization parameter *γ* in Eq. (1) has been adjusted to 1 through two-fold cross-validation, and we have chosen a forgetting factor of λ = 0.975 for capturing the data dynamics resulting in an *effective* data length of 10 s, long enough to ensure estimation stability.

As for the encoder model, we have used a Gaussian dictionary **G**_0_ to enforce smoothness in the TRF estimates. The columns of **G**_0_ consist of overlapping Gaussian kernels with the standard deviation of 20 ms whose means cover the 0s to 0.4s lag range with *T_s_* = 5 ms increments. The 20 ms standard deviation is consistent with the average full width at half maximum (FWHM) of an auditory MEG evoked response (M50 or M100), empirically obtained from MEG studies [17]. Thus, the overall dictionary discussed in Remark 2 takes the form **G** = diag (1, **G**_0_, **G**_0_). Also, similar to [17], we have used the logarithm of the speech envelopes as the regression covariates. Finally, the parameters of the FASTA package in encoder estimation have been chosen similar to those in the foregoing EEG analysis.

The M100 component of the TRF has shown to be more significant for the attended speaker than the unattended speaker [13, 17]. Thus, at each instance *k*, we extract the magnitude of the negative peak close to the 0.1 s delay in the real-time TRF estimate of each speaker as the attention markers 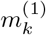 and 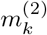. For the state-space model and the fixed-lag window, we have used the same configuration as in our foregoing EEG analysis, i.e. *K_W_* = ⌊15*f_s_*/*W*⌋, *K_F_* = ⌊1.5*f_s_*/*W*⌋, *a*_0_ = 2.008, and *b*_0_ = 0.2016. Note that the total delay in estimating the attentional state is now only 1.5 s, given that we use an encoding model for our MEG analysis. Furthermore, the prior distribution parameters for each subject were chosen according to the two fitted Log-Normal distributions on the extracted M100 values in the first 15 s of the trials, while choosing large variances for the Gamma priors to be non-informative. Similar to the preceding cases, the first 15 s of each trial can be thought of as an initialization stage.

#### 3.3.2. Estimation Results

Figure 11 shows our estimation results for four sample trials from the constant-attention (cases 1 and 2) and attention-switch (cases 3 and 4) experiments. For graphical convenience, we have rearranged the MEG data such that in the constant-attention experiment, the attention is always on speaker 1, and in the attention-switch experiment, speaker 1 is attended from 0s to 28 s. Cases 1 and 3 corresponds to trials in which the extracted M100 values for the attended speaker are more significant than those of the unattended speaker during most of the trial duration. Cases 2 and 4, on the other hand, correspond to trials in which the extracted M100 values are not reliable representatives of the attentional state. Row A in Fig. 11 shows the estimated TRFs for speakers 1 and 2 in time for each of the four cases. The location of the M100 peaks is shown and tracked with a narrow line (yellow) on the extracted M100 components (blue). The M50 components are also evident as positive peaks occurring around the 50 ms lag. The M50 components do not strongly depend on the attentional state of the listener [17, 13, 35, 36], which is consistent with those shown in Fig. 11-A.

**Figure 11:**
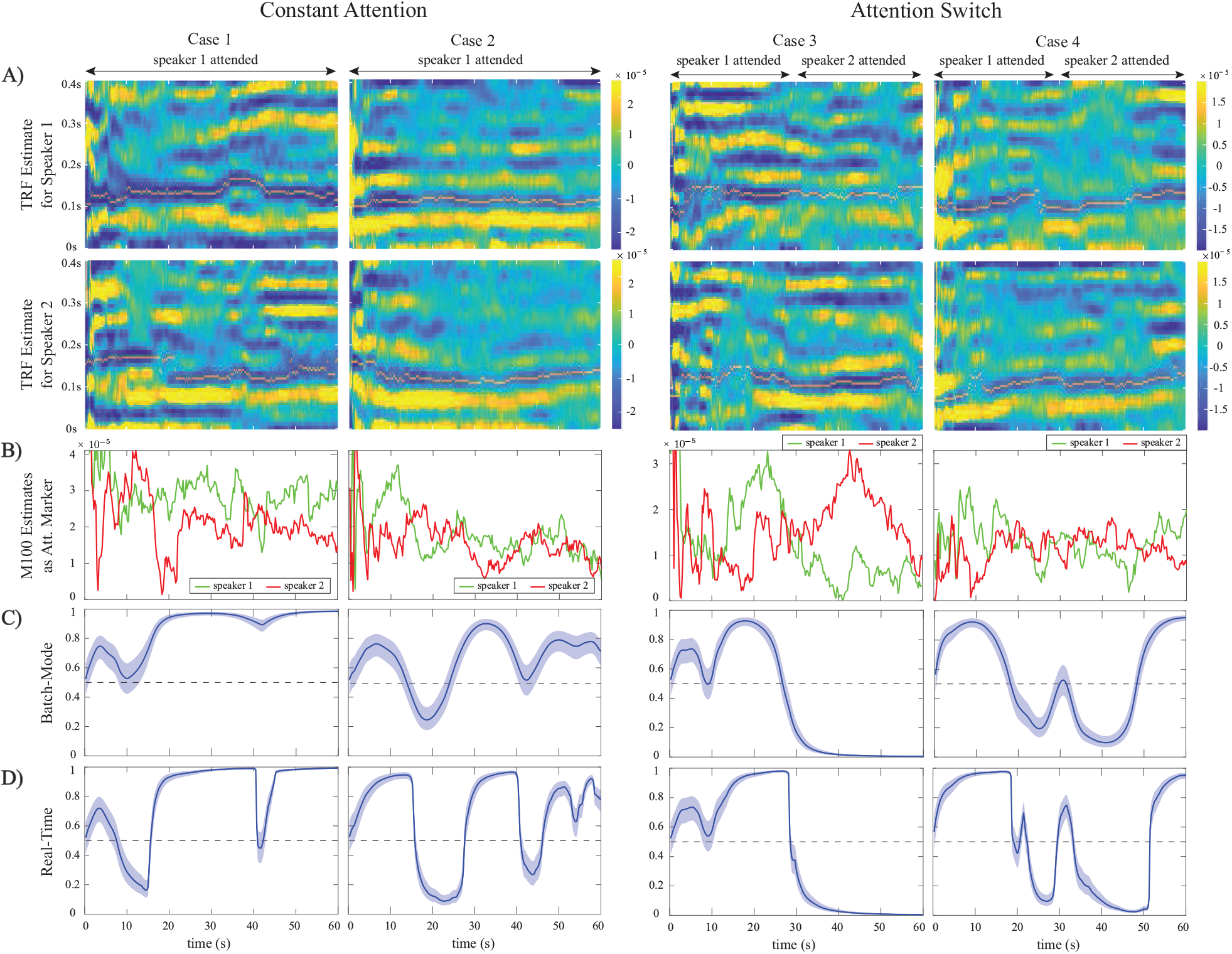
Examples from the constant-attention and attention-switch MEG experiments, using the M100 attention marker, for trials with reliable (cases 1 and 3) and unreliable (cases 2 and 4) separation of the attended and unattended speakers. A) TRF estimates for speakers 1 and 2 over time with the extracted M100 peak positions tracked by a narrow yellow line. B) Extracted M100 peak magnitudes over time for speakers 1 and 2 as the attention marker. In cases 1 and 3, the M100 components exhibit a strong modulation effect of the attentional state, i.e., the attended speaker has a larger M100 peak, in contrast to cases 2 and 4, where there is a weak modulation. C) Batch-mode state-space estimates of the attentional state. D) Real-time state-space estimates of the attentional state. The strong or weak modulation effects of attentional state in the extracted M100 components directly affects the classification accuracy and the width of the confidence intervals for both the batch-mode and real-time estimators.

Row B in Fig. 11 displays the extracted M100 peak magnitudes over time for speakers 1 and 2. The attention modulation effect is more significant in cases 1 and 3. Rows C and D respectively show the batch-mode and real-time estimates of the attentional state based on the extracted M100 values. As expected, the batch-mode output is more robust to the fluctuations in the extracted M100 peak values, with smoother transitions and larger confidence intervals. Despite the poor attention modulation effect in cases 2 and 4, we observe that both the real-time and the batch-mode state-space models show reasonable performance in translating the extracted M100 peak values to a robust measure of the attentional state. This effect is notable in Rows C and D of Case 4. We performed the same analysis as in Fig. 9 to assess the effect of the forward-lag parameter *K_F_*. Since the results were quite similar to those in Figures 6 and 9, we have omitted them for brevity and chose the same forward-lag of 1.5 s.

Finally, Fig. 12 summarizes the *real-time* classification results for the constant-attention (left panels) and attention-switch (right panels) MEG experiments. The classification convention is similar to that used in our EEG analysis, and is illustrated in Fig. 12-A for the completeness. For the attention-switch experiment, the 28 s-30 s interval is removed from the classification analysis, as it pertains to a silence period during which the subject is instructed to switch attention. Fig. 12-B shows the corresponding classification results, consisting of 36 trials for the constant-attention and 18 trials for the attention-switch experiments. Each circle corresponds to a single trial and the subjects in each experiment are color-coded. The average classification results per trial are shown in Fig. 12-C for each subject. The average hit rate and false alarm rates in the constant-attention experiments are respectively given by 71.67% and 20.81%. These quantities for the attention-switch experiment are respectively given by 64.12% and 26.16%, showing a reduction in hit rate and increase in false alarm.

**Figure 12:**
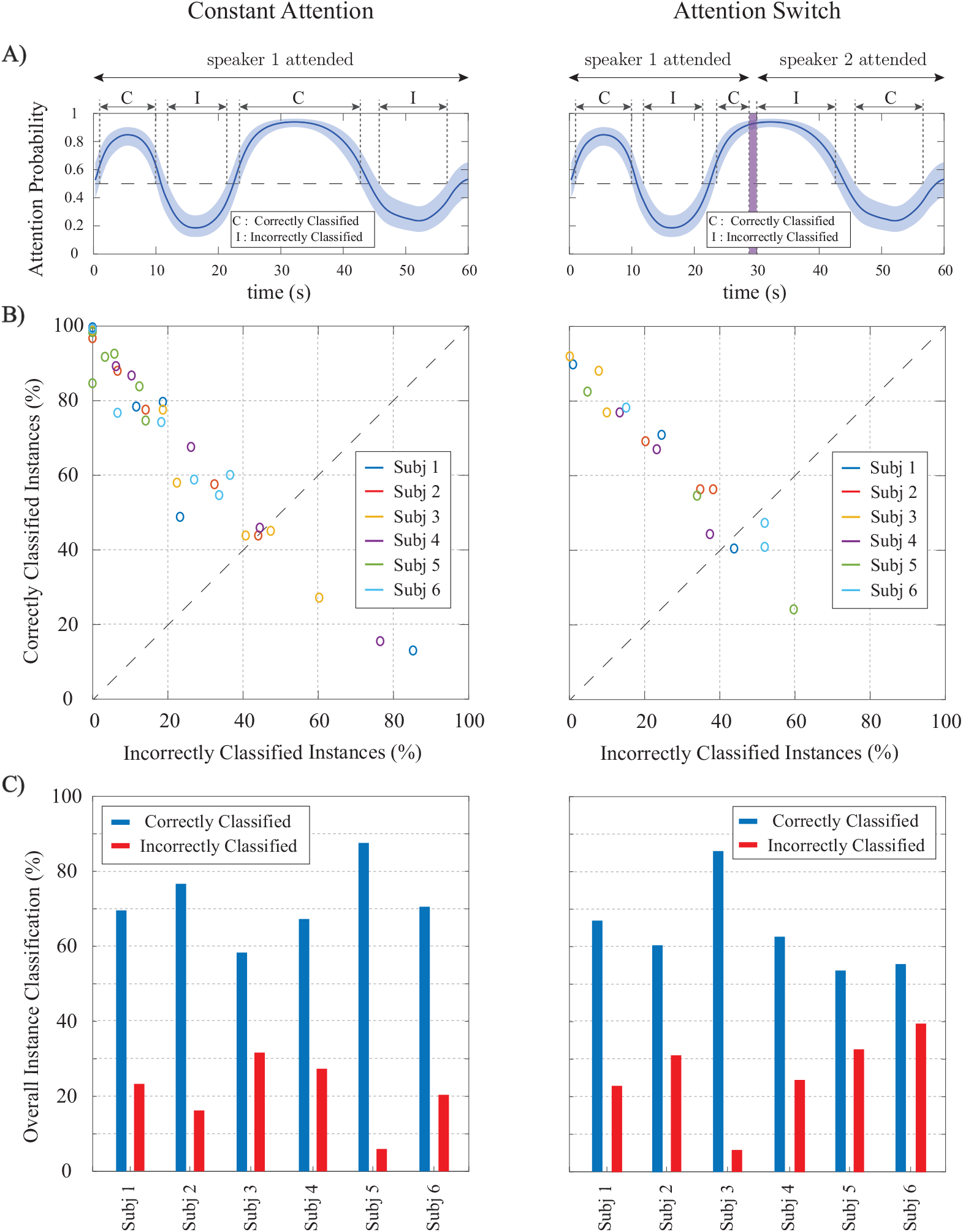
Summary of real-time classification results for the constant-attention (left panels) and attention-switch (right panels) MEG experiments. A) a generic instance of the state-space output for a trial illustrating the classification convention. B) Classification results per trial for all subjects; each circle corresponds to a trial and the subjects are color-coded. The trials falling below the dashed line have more incorrectly classified instances than correctly classified ones. C) Average classification performance over all trials for the six subjects.

## 4. Discussion

In this work, we have proposed a framework for real-time decoding of the attentional state of a listener in a dual-speaker environment from M/EEG. This framework consists of three modules. In the first module, the encoding/decoding coefficients, relating the neural response to the envelopes of the two speech streams, are estimated in a low-complexity and real-time fashion. Existing approaches for encoder/decoder estimation operate in an offline fashion using multiple experiment trials or large training datasets [10, 16, 28, 25], and hence are not suitable for real-time applications. To address this issue, we have integrated the forgetting factor mechanism used in adaptive filtering with ℓ_1_-regularization, in order to capture the coefficient dynamics and mitigate overfitting.

In the second module, a function of the estimated encoding/decoding coefficients and the acoustic data, which we refer to as the *attention marker*, is calculated in real-time for each speaker. The role of the attention marker is to provide dynamic features that create statistical separation between the attended and the unattended speakers. Examples of such attention markers include correlation-based measures (e.g. correlation of the acoustic envelopes and their reconstruction from neural response), or measures solely based on the estimated decoding/encoding coefficients (e.g. the ℓ_1_-norm of the decoder coefficients or the M100 peak of the encoder).

Finally, the attention marker is passed to the third module consisting of a near real-time state-space estimator. To control the delay in state estimation, we adopt a fixed-lag smoothing paradigm, in which the past and near future data are used to estimate the states. The role of the state-space model is to translate the noisy and highly variable attention markers to robust measures of the attentional state with minimal delay. We have archived a publicly available MATLAB implementation of our framework on the open source repository GitHub in order to ease reproducibility [37].

We validated the performance of our proposed framework using simulated EEG and MEG data, in which the ground truth attentional states are known. We also applied our proposed methods to experimentally recorded MEG and EEG data. As for a comparison benchmark, we considered the offline state-space attention decoding approach of [16]. Our MEG analysis showed that although the proposed real-time estimator has access to significantly fewer data points, it closely matches the outcome of the offline state-space estimator in [16], for which the entire data from multiple trials are used for attention decoding. In particular, our analysis of the MEG data in constant-attention conditions revealed a hit rate of ~ 70% and a false alarm rate of ~ 20% at the group level. While the performance is slightly degraded compared to the offline analysis of [16], our algorithms operate in real-time with 1.5s forward delay, over single trials, and using minimal tuning. Similarly, our analysis of EEG data provided ~ 80% hit rate and ~ 15% false alarm rate at a single trial level. These performance measures are slightly degraded compared to the results of offline approaches such as [10].

Our proposed modular design admits the use of any attention-modulated statistic or feature as the attention marker, three of which have been considered in this work. While some attention markers perform better than the rest in certain applications, our goal in this work was to provide different examples of attention markers which can be used in the encoding/decoding models based on the literature, rather than comparing their performance against each other. The choice of the best attention marker that results in the highest classification accuracy is a problem-specific matter. Our modular design allows to evaluate the performance of a variety of attention markers for a given experimental setting, while fixing the encoding/decoding estimation and state-space modules, and to choose one that provides the desired classification performance.

A practical limitation of our proposed methodology in its current form is the need to have access to clean acoustic data in order to form regressors based on the speech envelopes. In a realistic scenario, the speaker envelopes have to be extracted from the noisy mixture of speeches recorded by microphone arrays. Thanks to a number of fairly recent results in attention decoding literature [28, 24, 26, 25, 27], it is possible to integrate our methodology with a pre-processing module that extracts the acoustic features of individual speech streams from their noisy mixtures. We view this extension as a future direction of research.

Our proposed framework has several advantages over existing methodologies. First, our algorithms require minimal amount of offline tuning or training. The subject-specific hyperparameters used by the algorithms are tuned prior to real-time application in a supervised manner. The only major offline tuning step in our framework is computing the subject-specific channel weights in the encoding model for MEG analysis in order to extract the auditory component of the neural response. This is due to the fact that the channel locations are not fixed with respect to the head position across subjects. It is worth noting that this step can be avoided if the encoding model treats the MEG channels separately in a multivariate model. Given that recent studies suggest that the M100 component of the encoder obtained from the MEG auditory response is a reliable attention marker [13, 14, 17], we adopted the DSS algorithm for computing the channel weights that compose the auditory response in an offline fashion.

Second, our analysis allows to characterize the performance of the attentional state classification using single trials, which is important for practical applications such as smart hearing aids. Existing studies based on offline algorithms perform classification based on cross-trial performance. For instance, in [10], for each 1 min of test trial, 29 mins of training data are used. In addition, the probabilistic output of our attentional state decoding framework can be used for further statistical analysis and soft-decision mechanisms which are desired in smart hearing aid applications. Finally, the modular design of our framework facilitates its adaptation to more complex auditory scenes (e.g. with multiple speakers and realistic noise and reverberation conditions) and integration of other covariates relevant to real-time applications (e.g. electrooculography measurements).

## Conflict of Interest Statement

The authors declare that the research was conducted in the absence of any commercial or financial relationships that could be construed as a potential conflict of interest.

## Author Contributions

TZ, JZS and BB designed the research. SM and BB performed the research, with contributions to the methods by AS, SA, and TZ and experimental data supplied by TZ and JZS. All authors participated in writing the paper.

## Funding

This material is based on work supported in part by the National Science Foundation Award No. 1552946 and a research gift from the Starkey Hearing Technologies to the University of Maryland.

## Acknowledgments

The authors would like to thank Dr. Tom Goldstein for helpful remarks on adapting the FASTA package options to our decoder/encoder estimation problem.

